## Supplementary figures and images for "Conformational dynamics and asymmetry in multimodal inhibition of membrane-bound pyrophosphatases"

### FigEV1

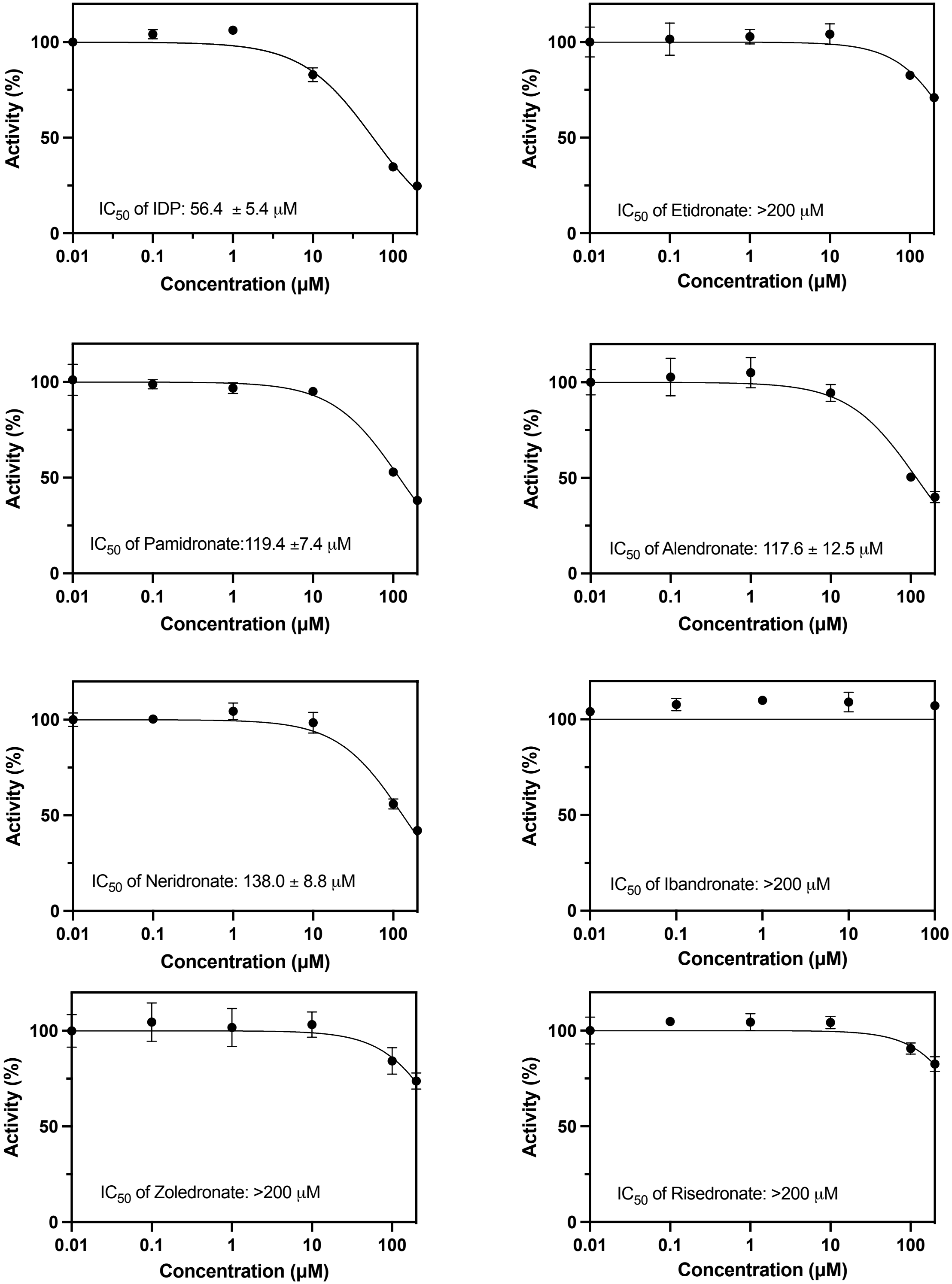

### FigEV2

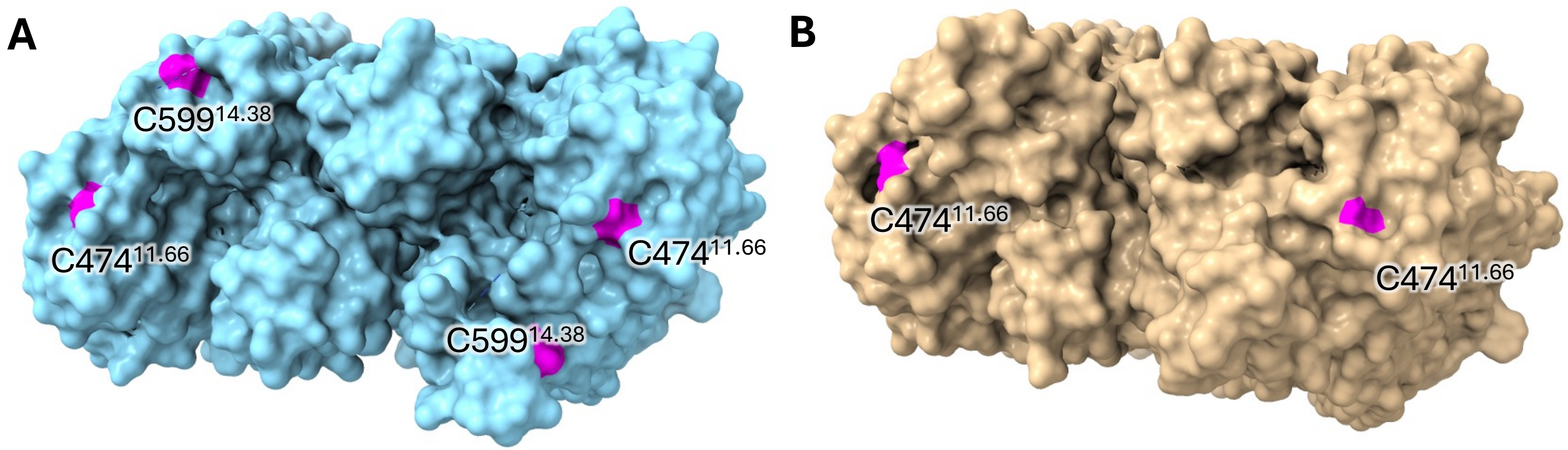

### FigEV3

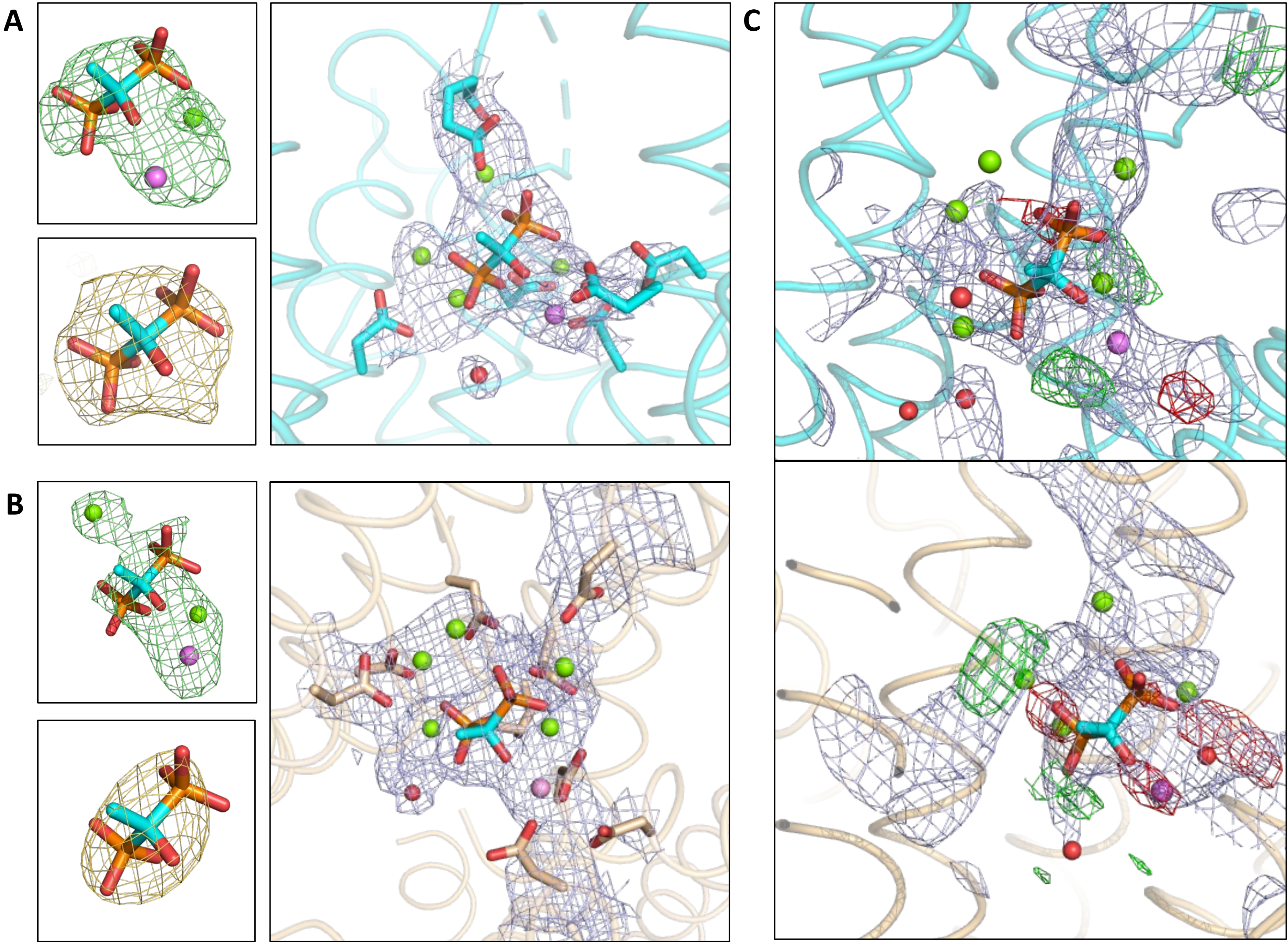

### FigEV4

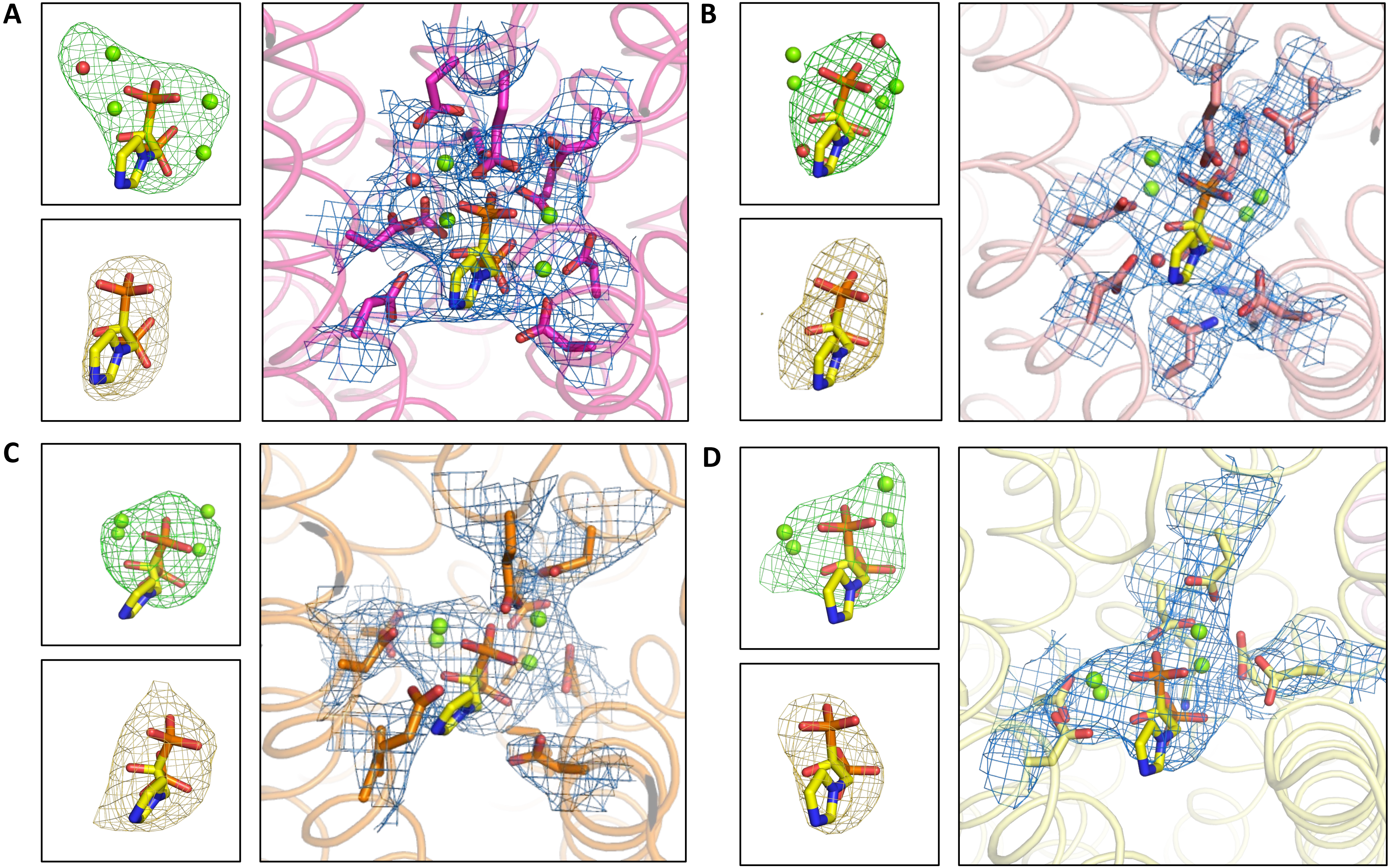

### FigEV5

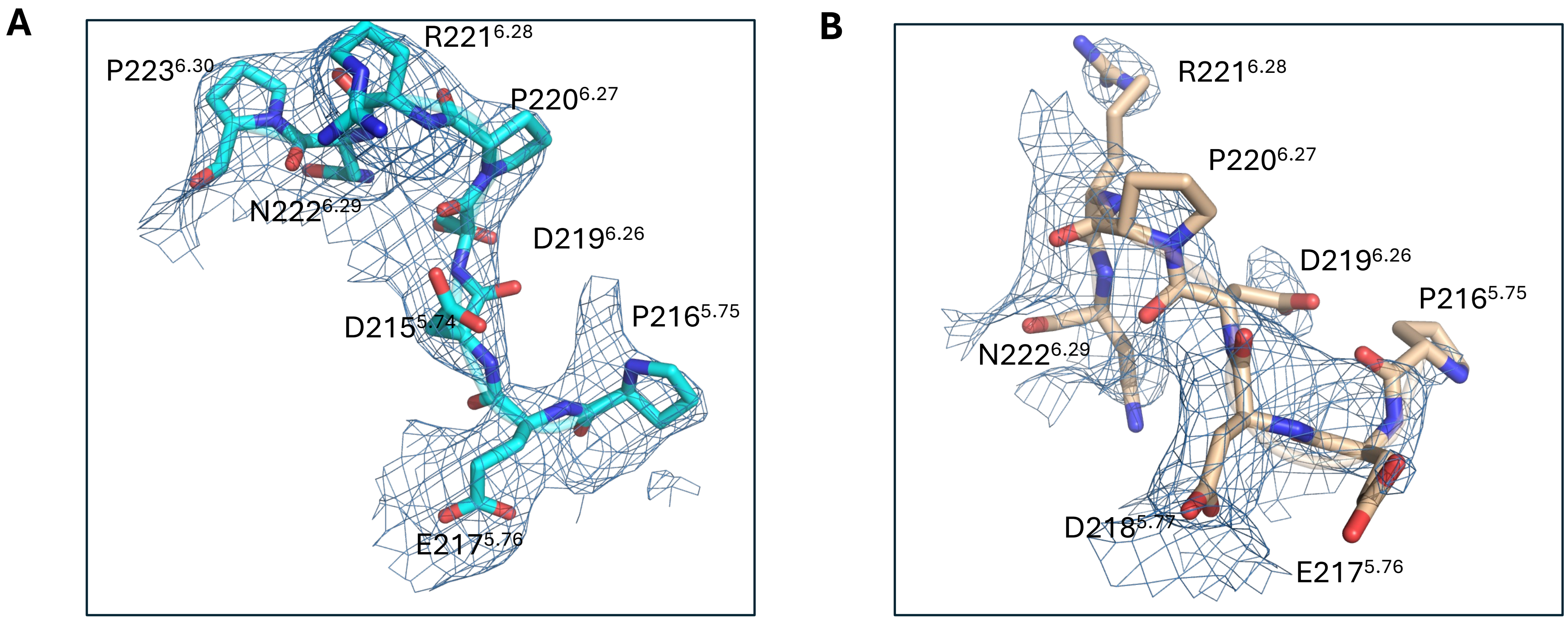

### FigEV6

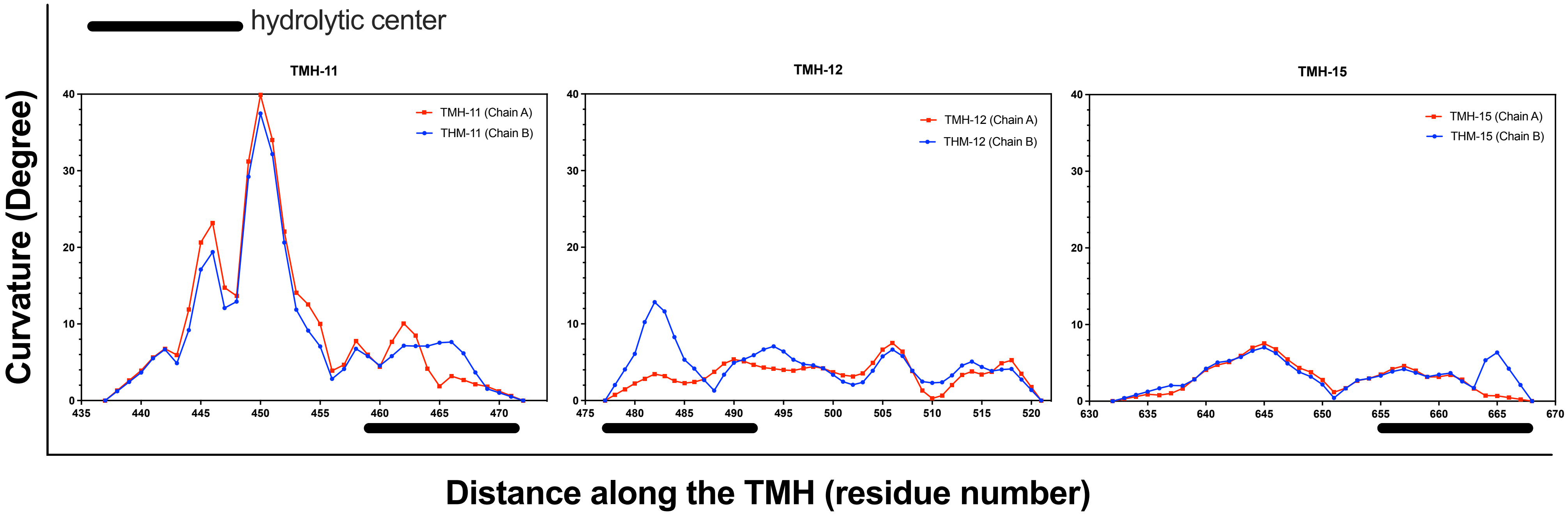

### FigEV7

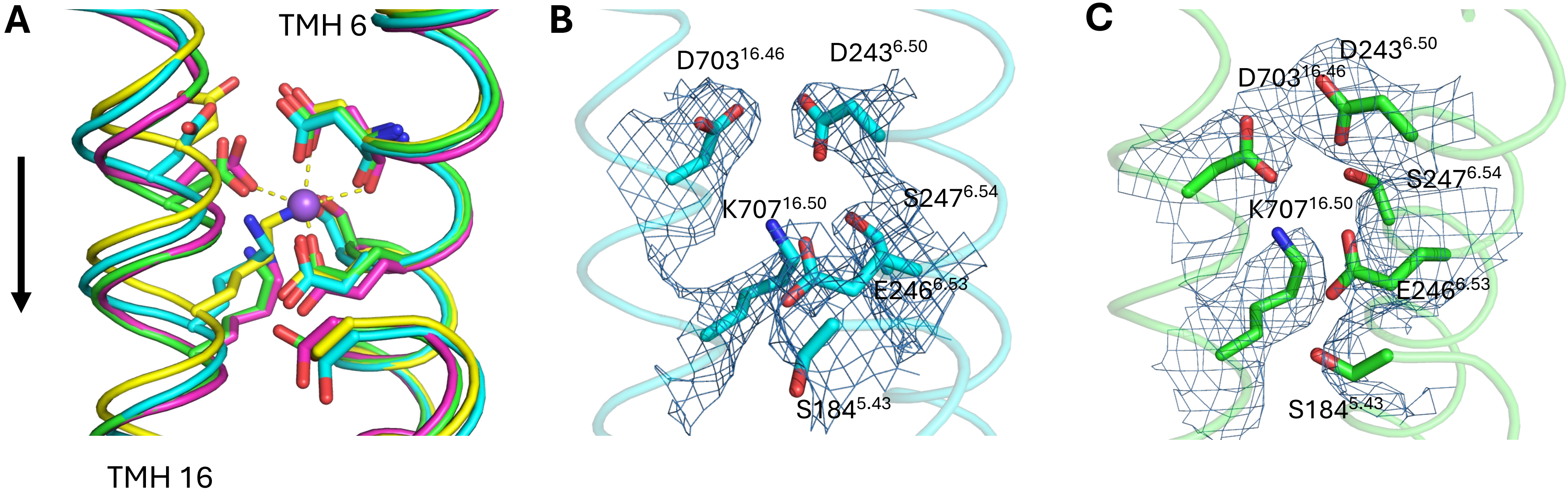

### FigEV8

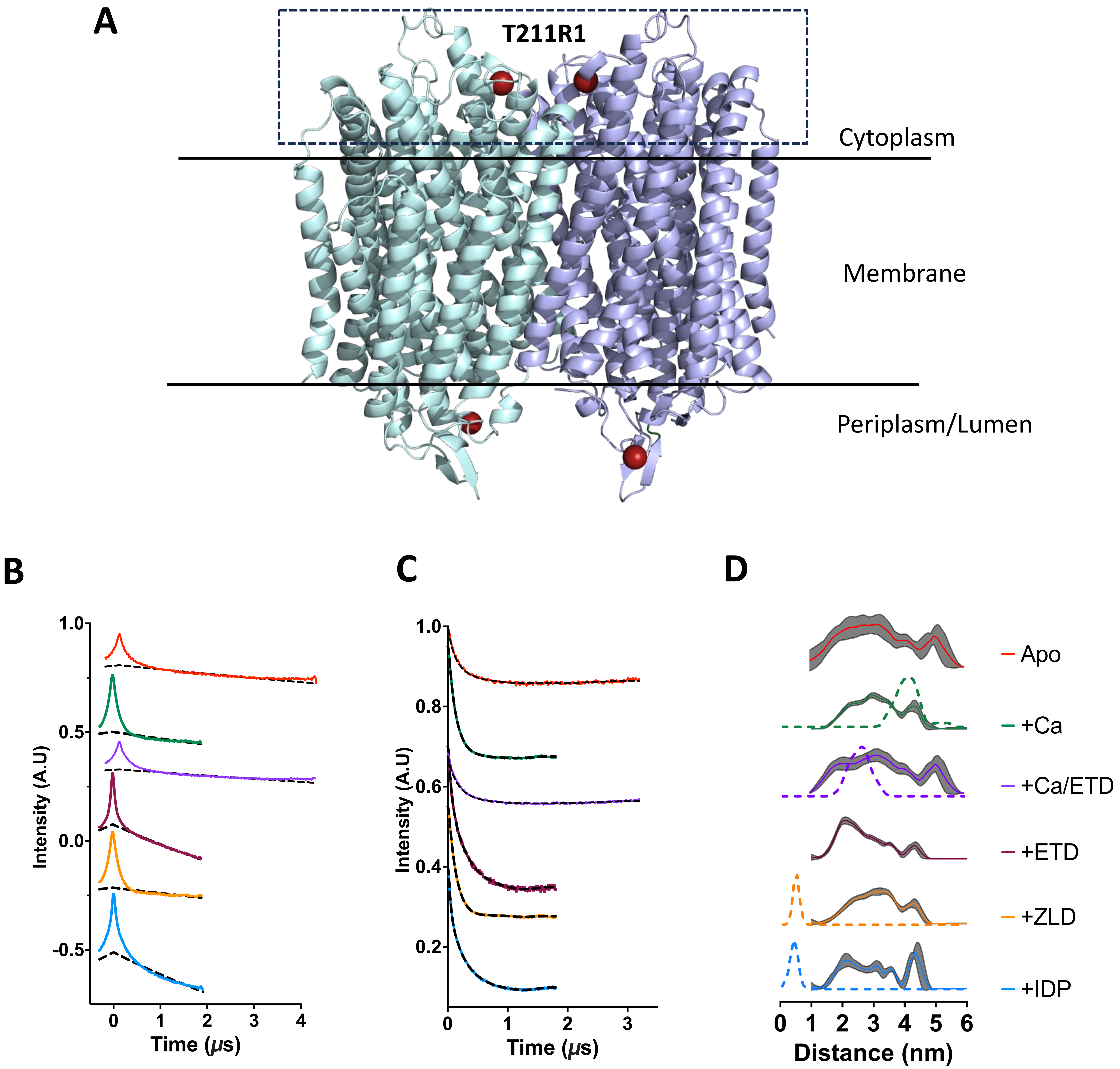

### FigEV9

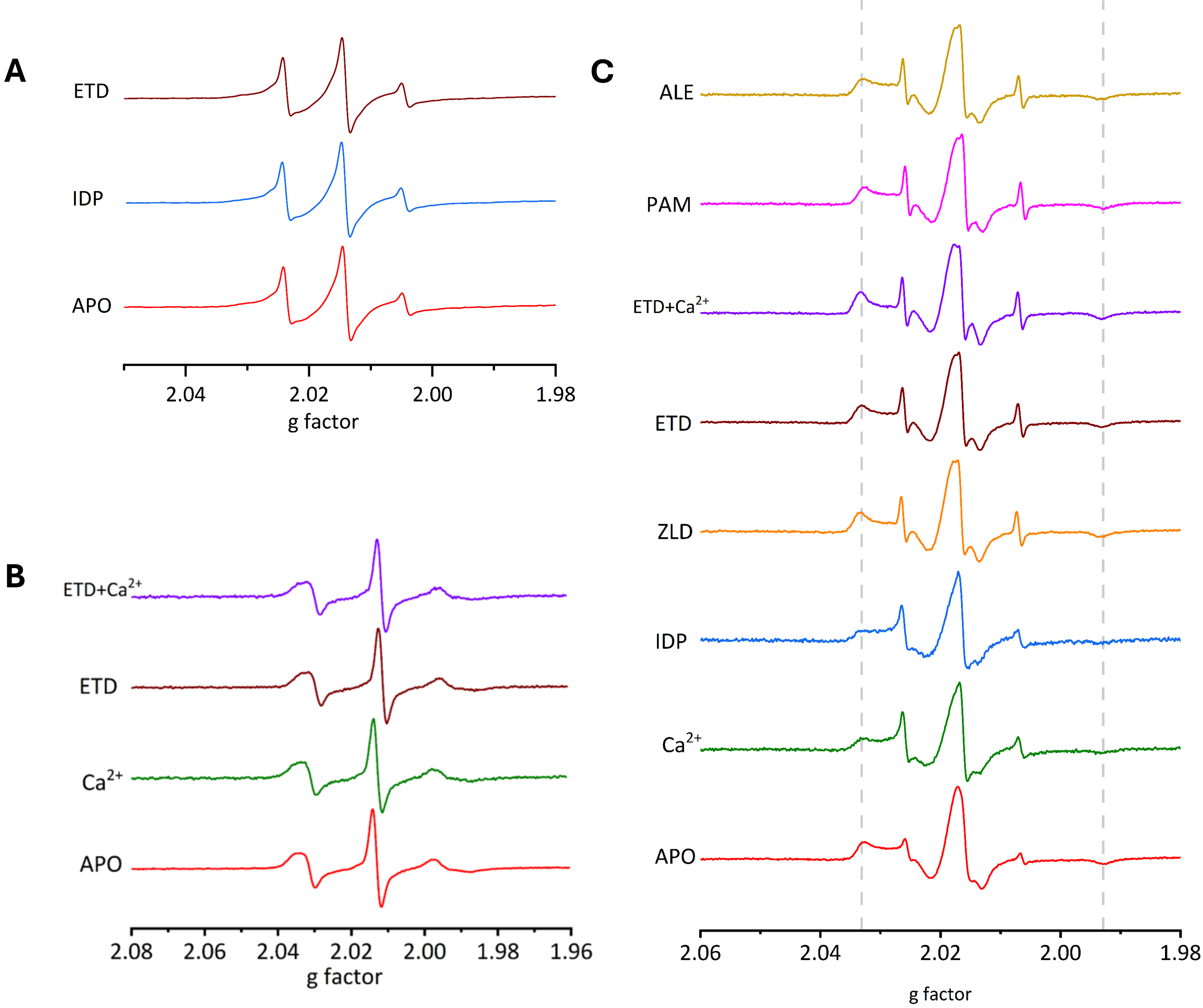

### FigEV10

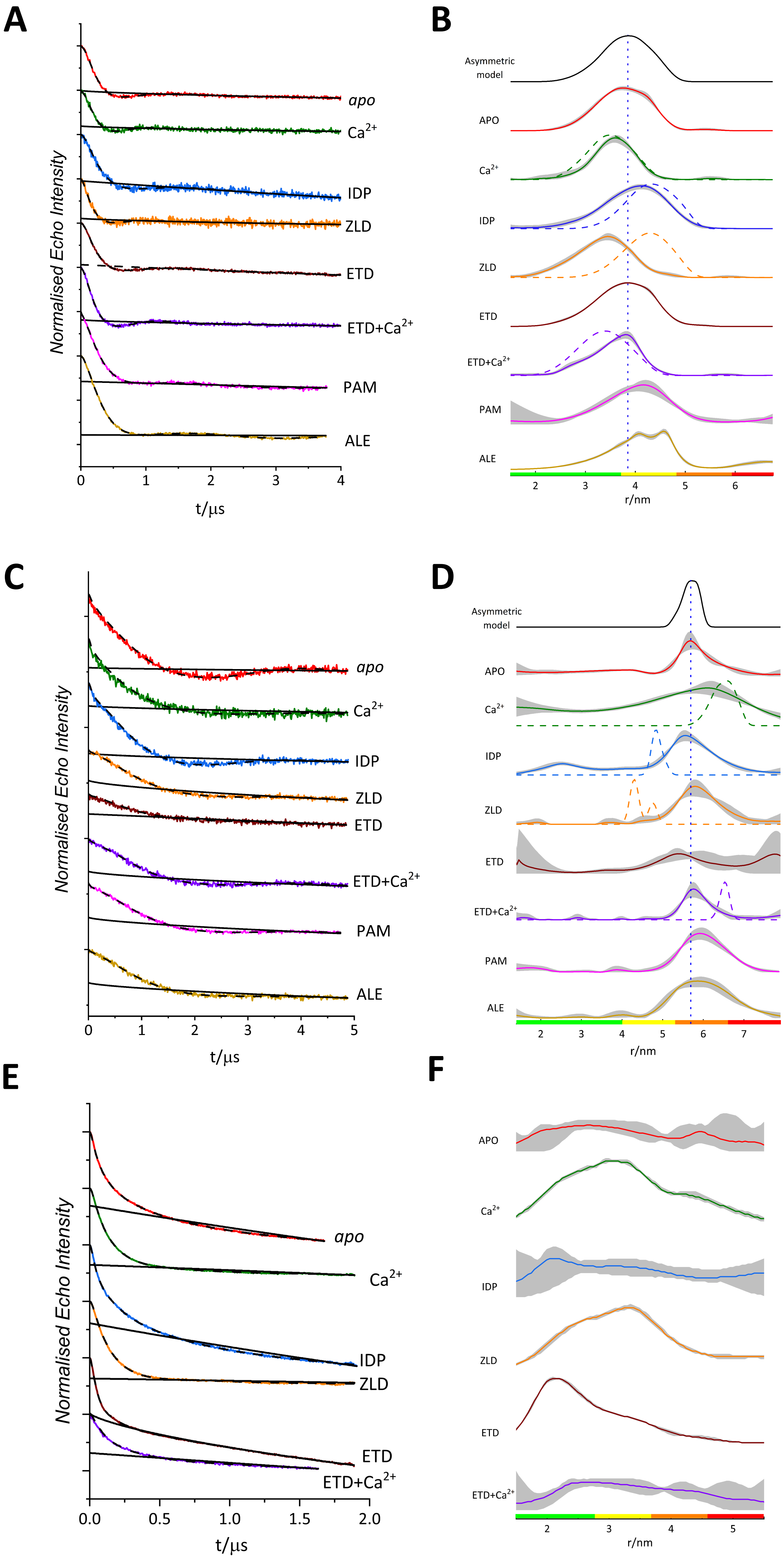
