## Supplementary material for "Conformational dynamics and asymmetry in multimodal inhibition of membrane-bound pyrophosphatases": FigEV11

### Slide 1
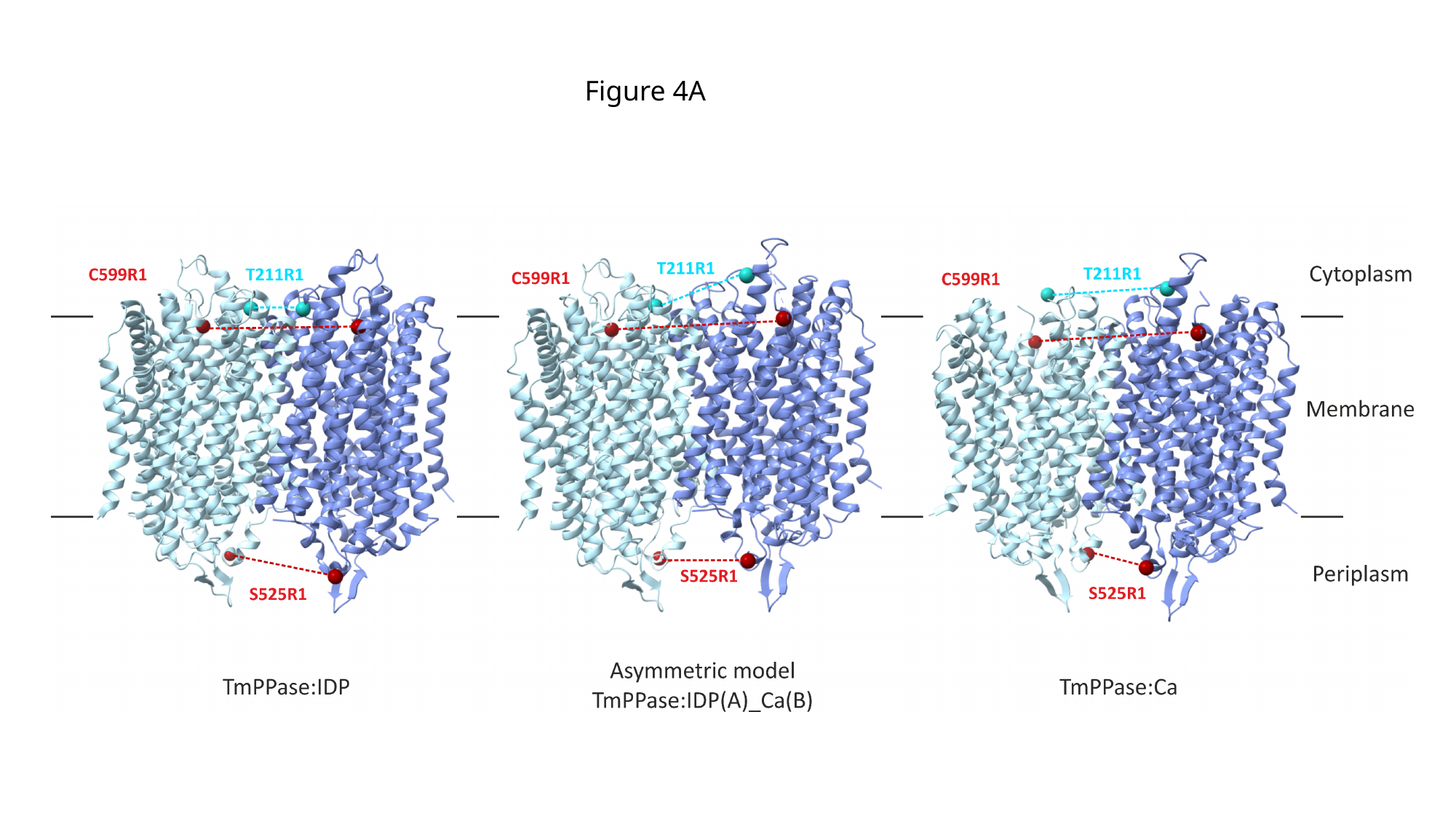

Figure 4A

### Slide 2
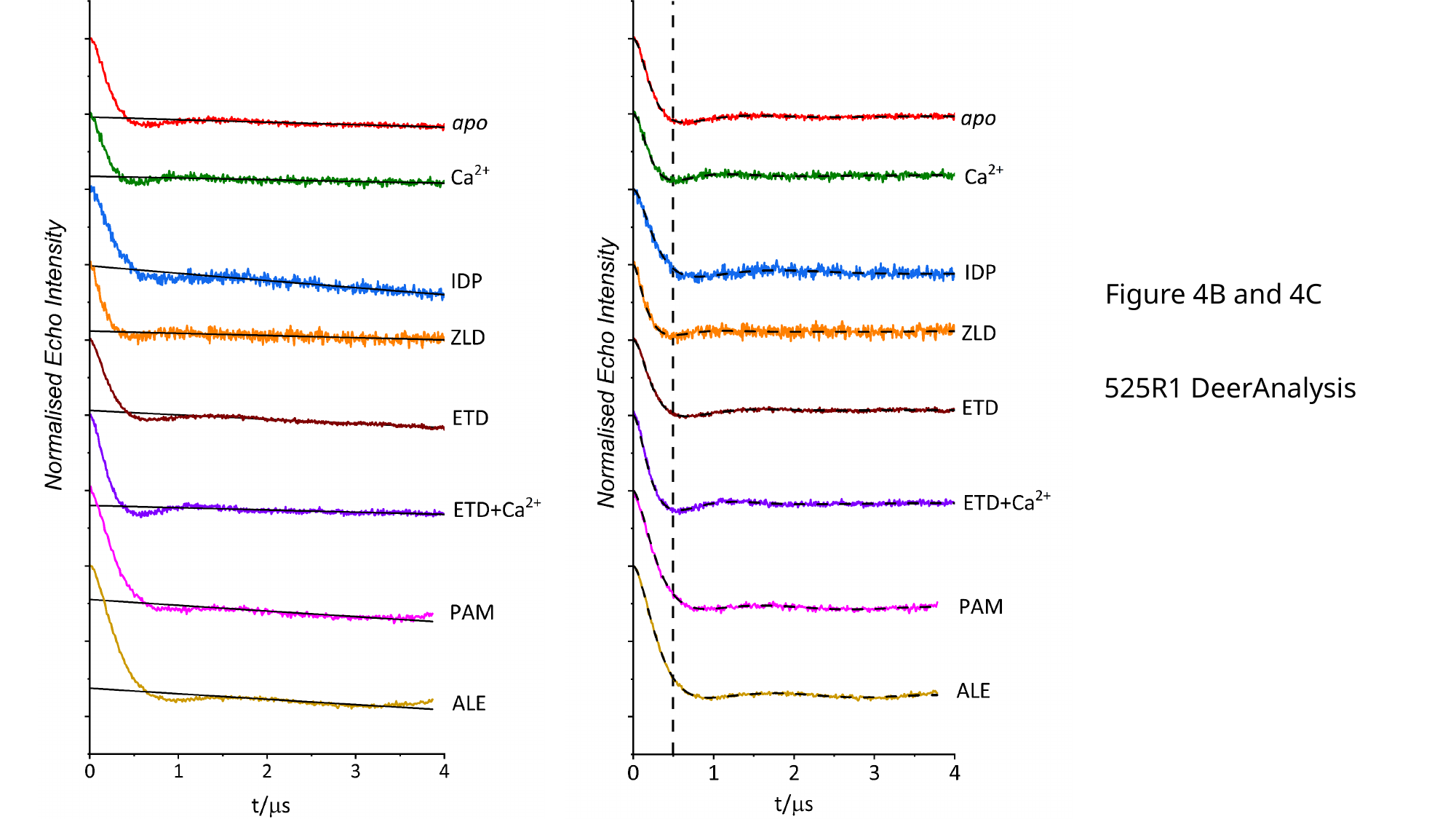

Figure 4B and 4C
525R1 DeerAnalysis

### Slide 3
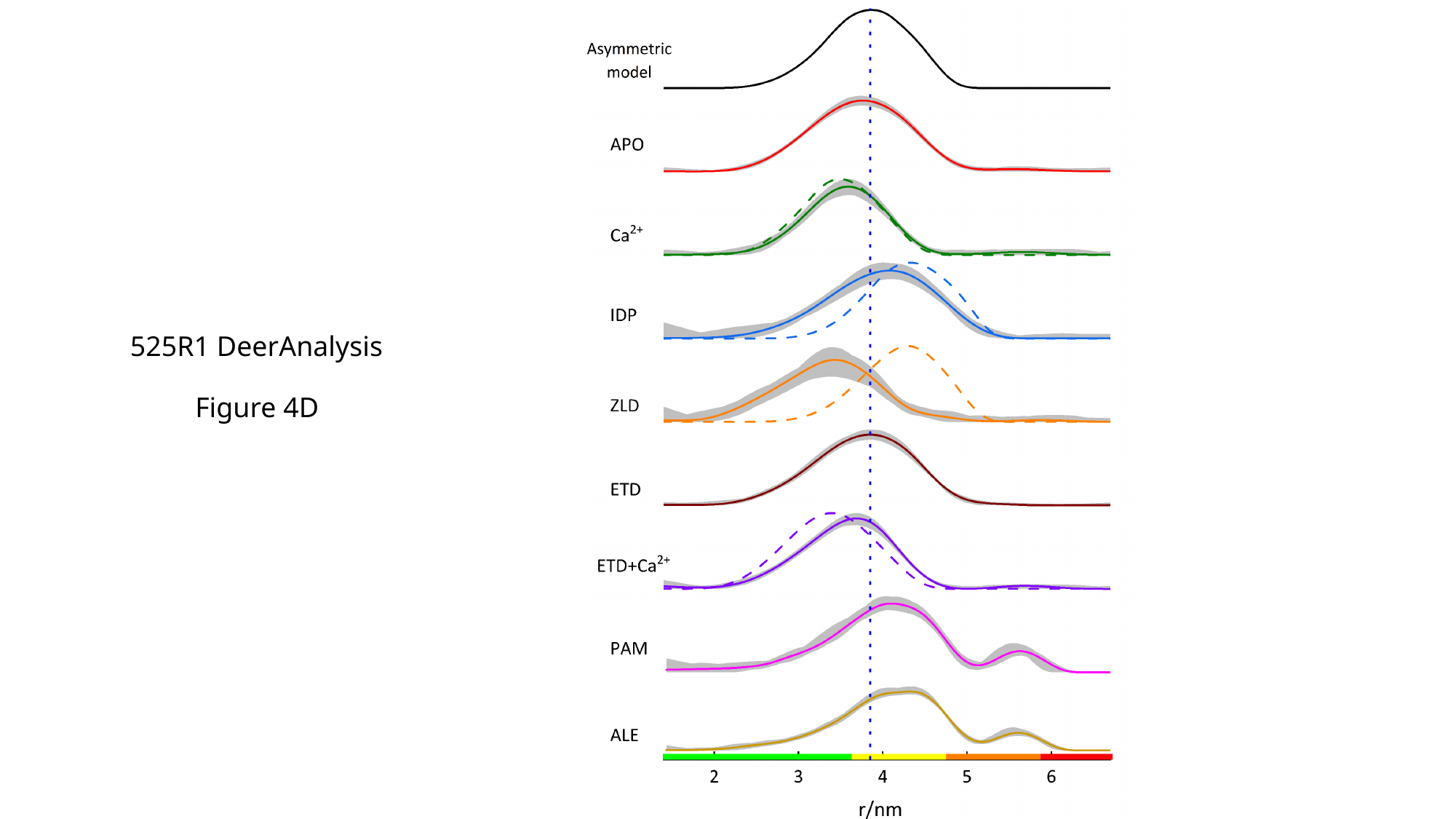

525R1 DeerAnalysis
Figure 4D

### Slide 4
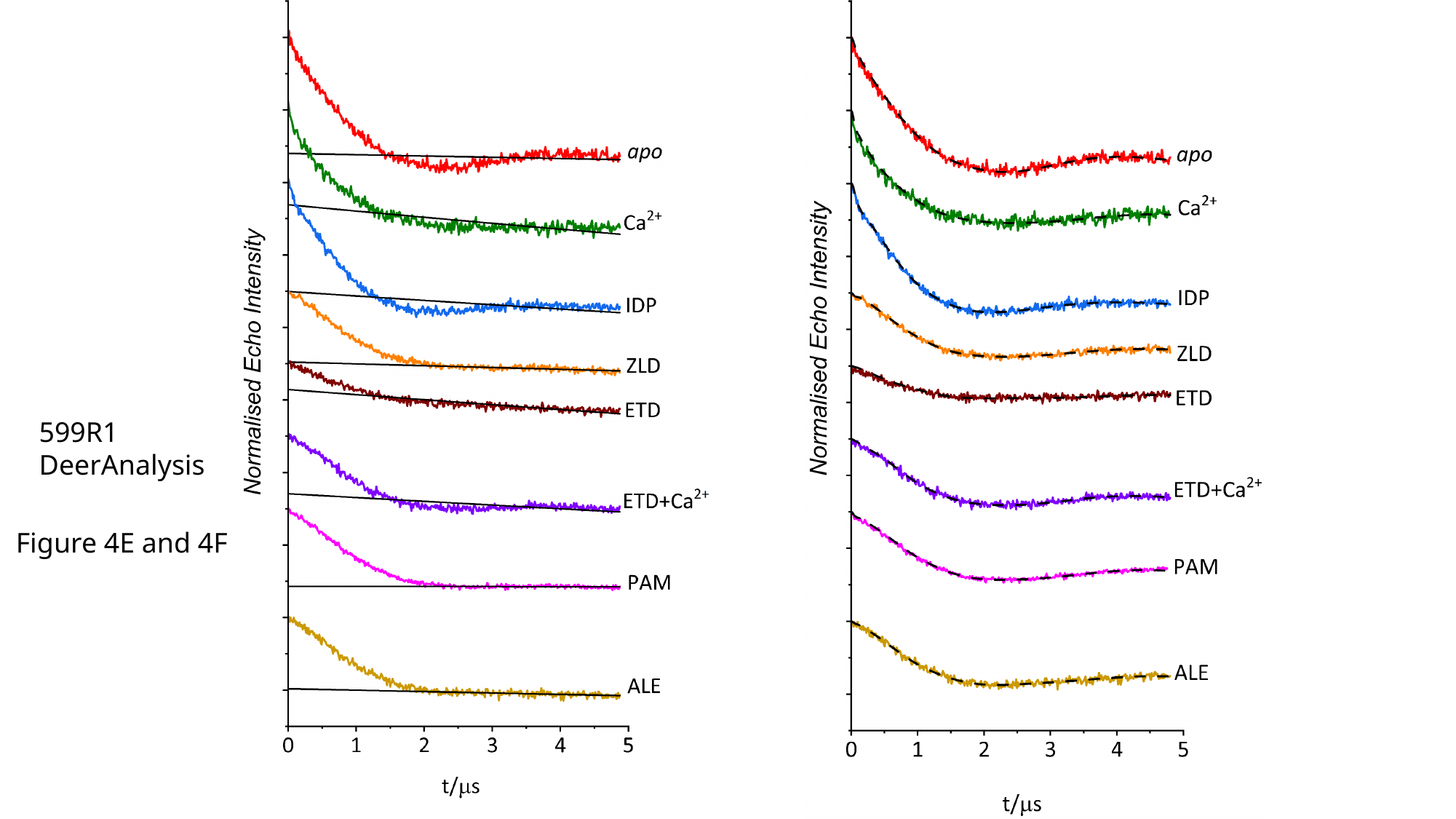

599R1
DeerAnalysis
Figure 4E and 4F

### Slide 5
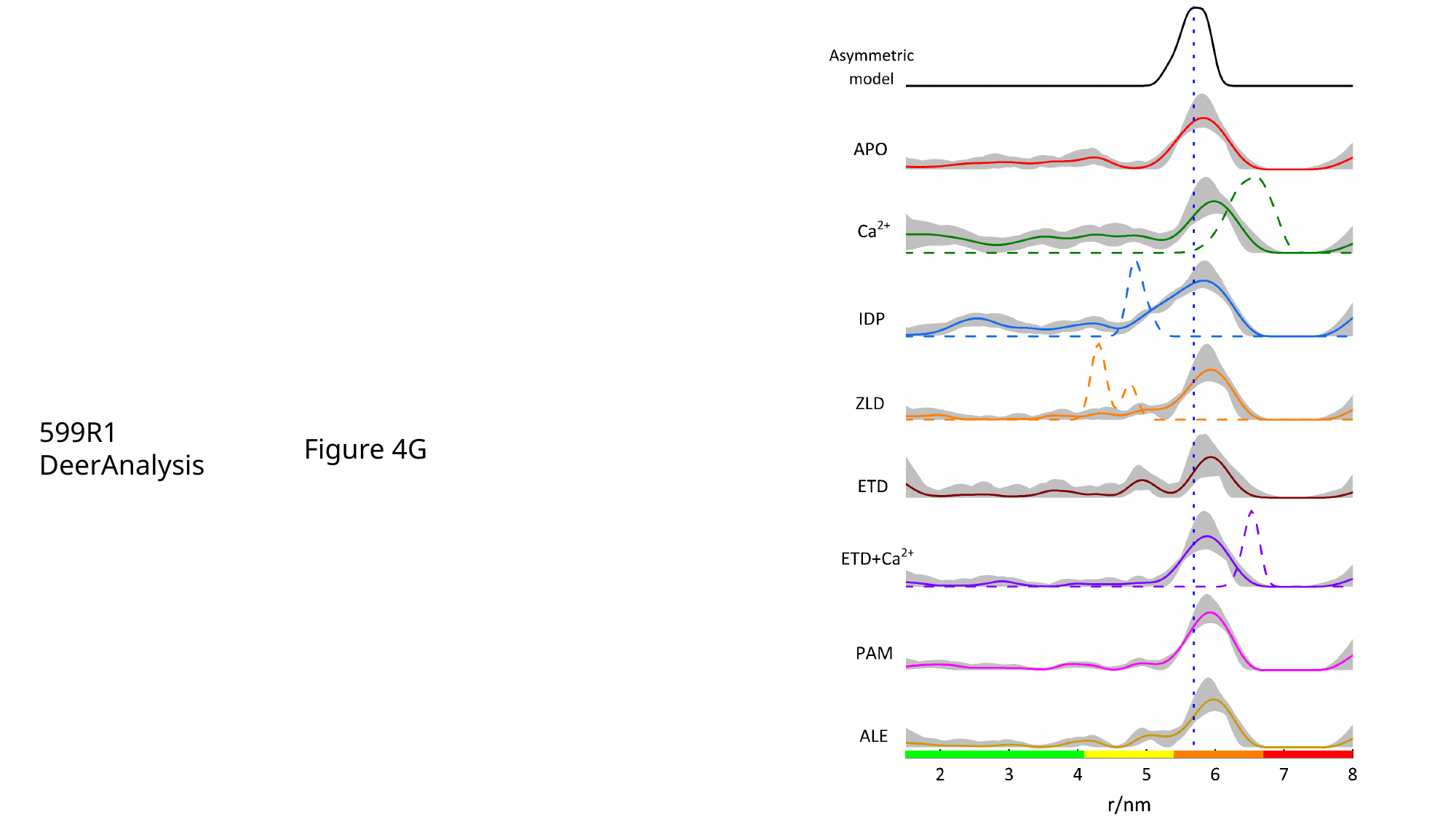

599R1
DeerAnalysis
Figure 4G

### Slide 6
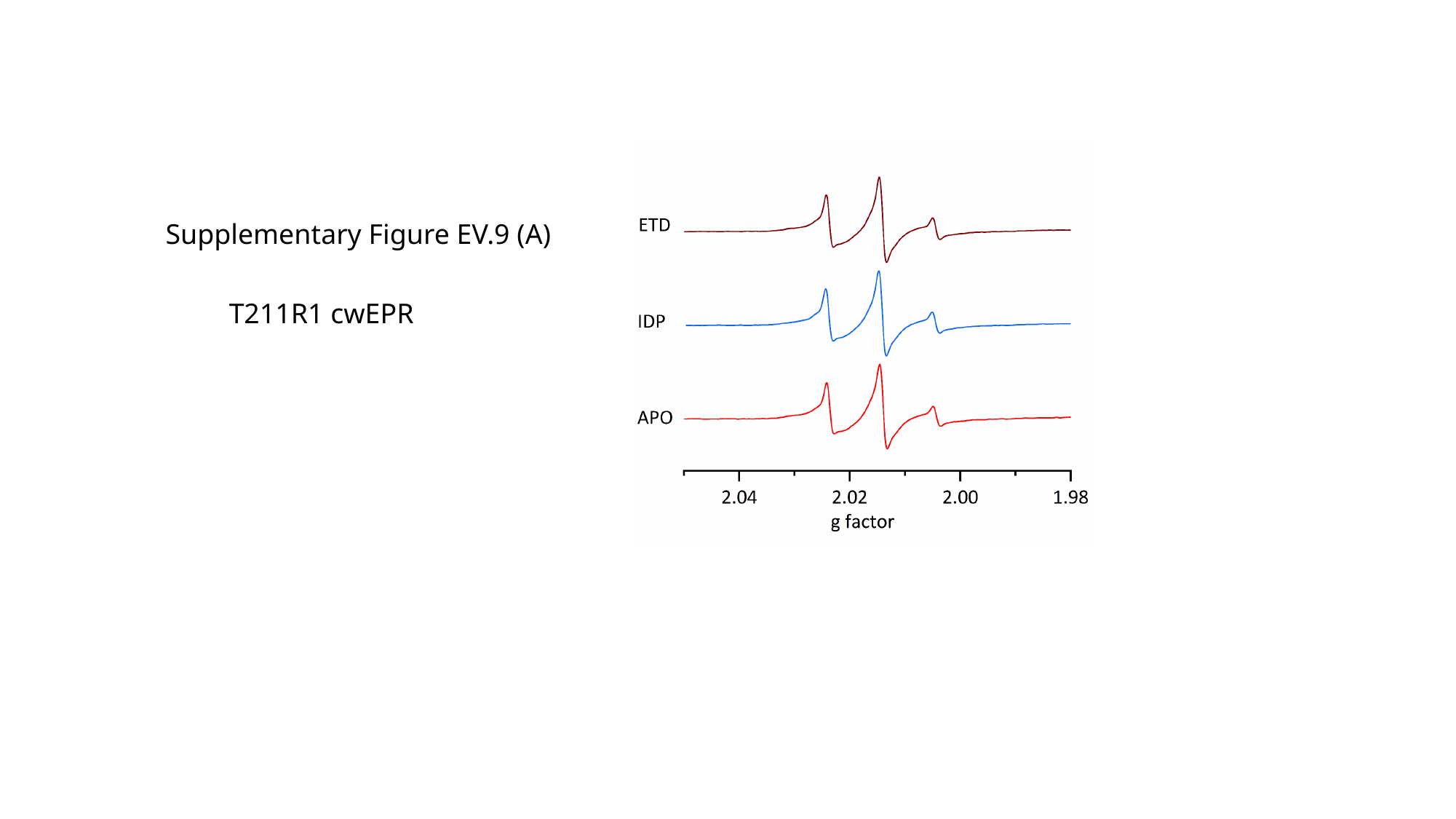

Supplementary Figure EV.9 (A)
T211R1 cwEPR

### Slide 7
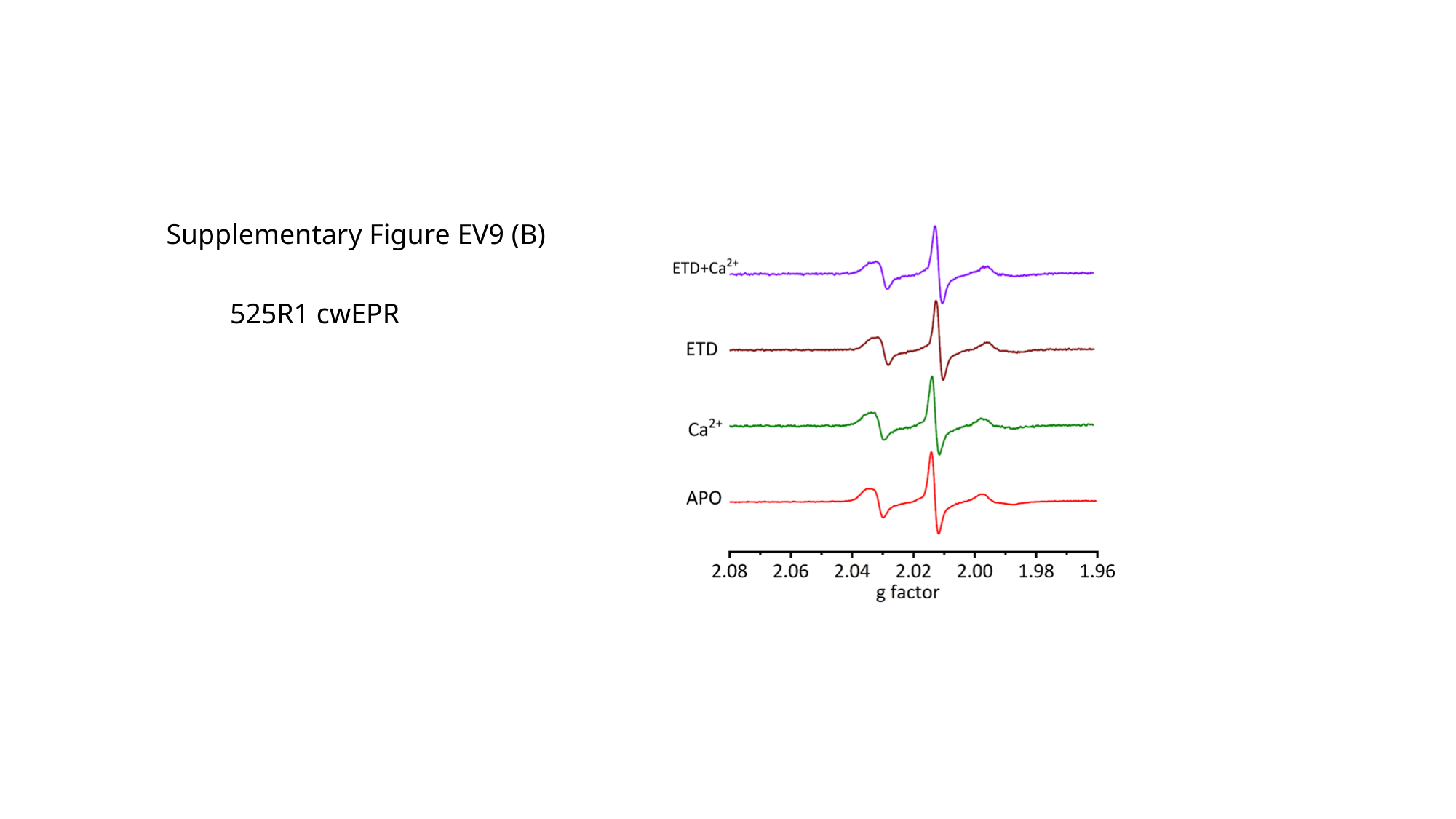

Supplementary Figure EV9 (B)
525R1 cwEPR

### Slide 8
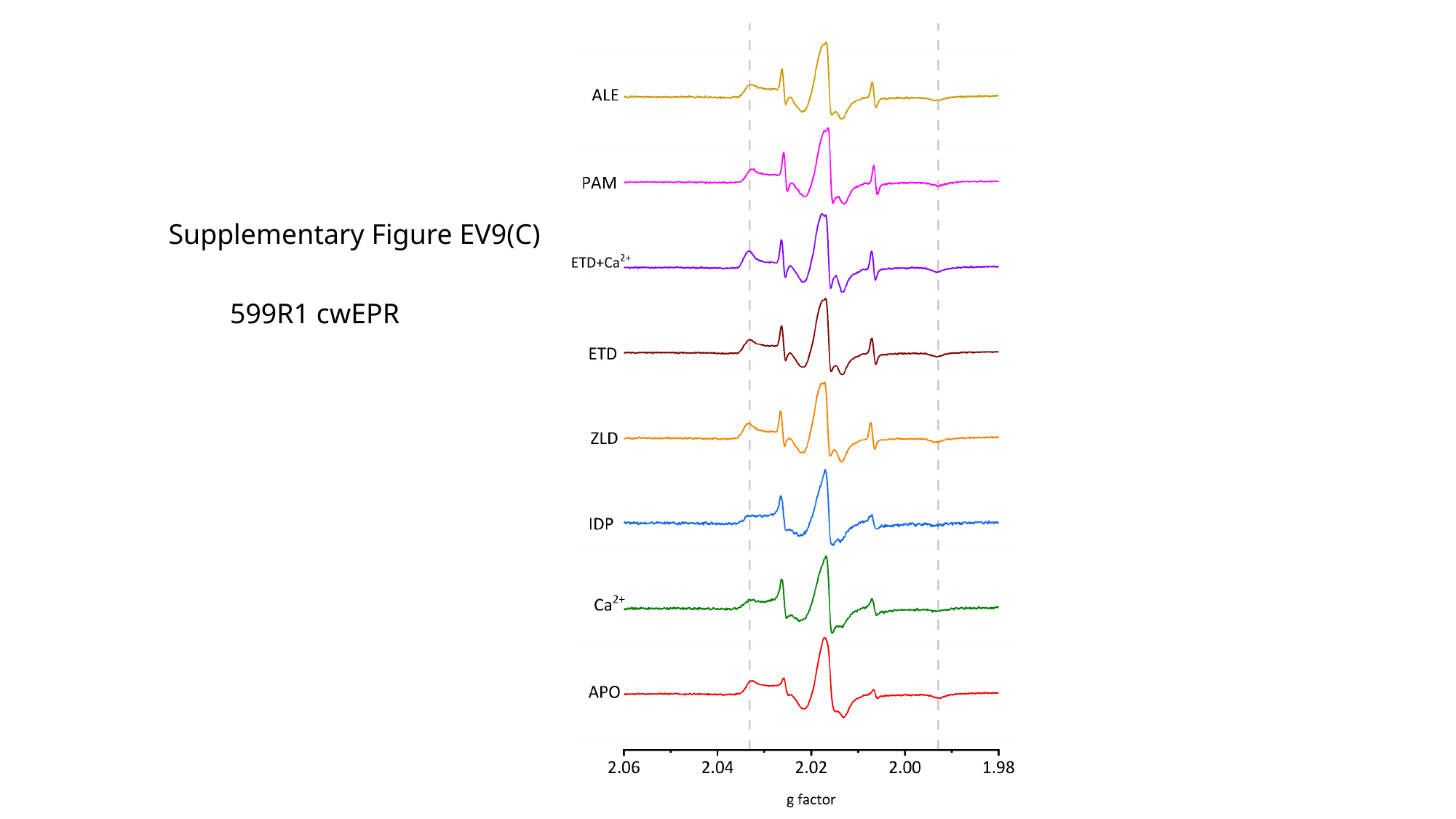

Supplementary Figure EV9(C)
599R1 cwEPR

### Slide 9
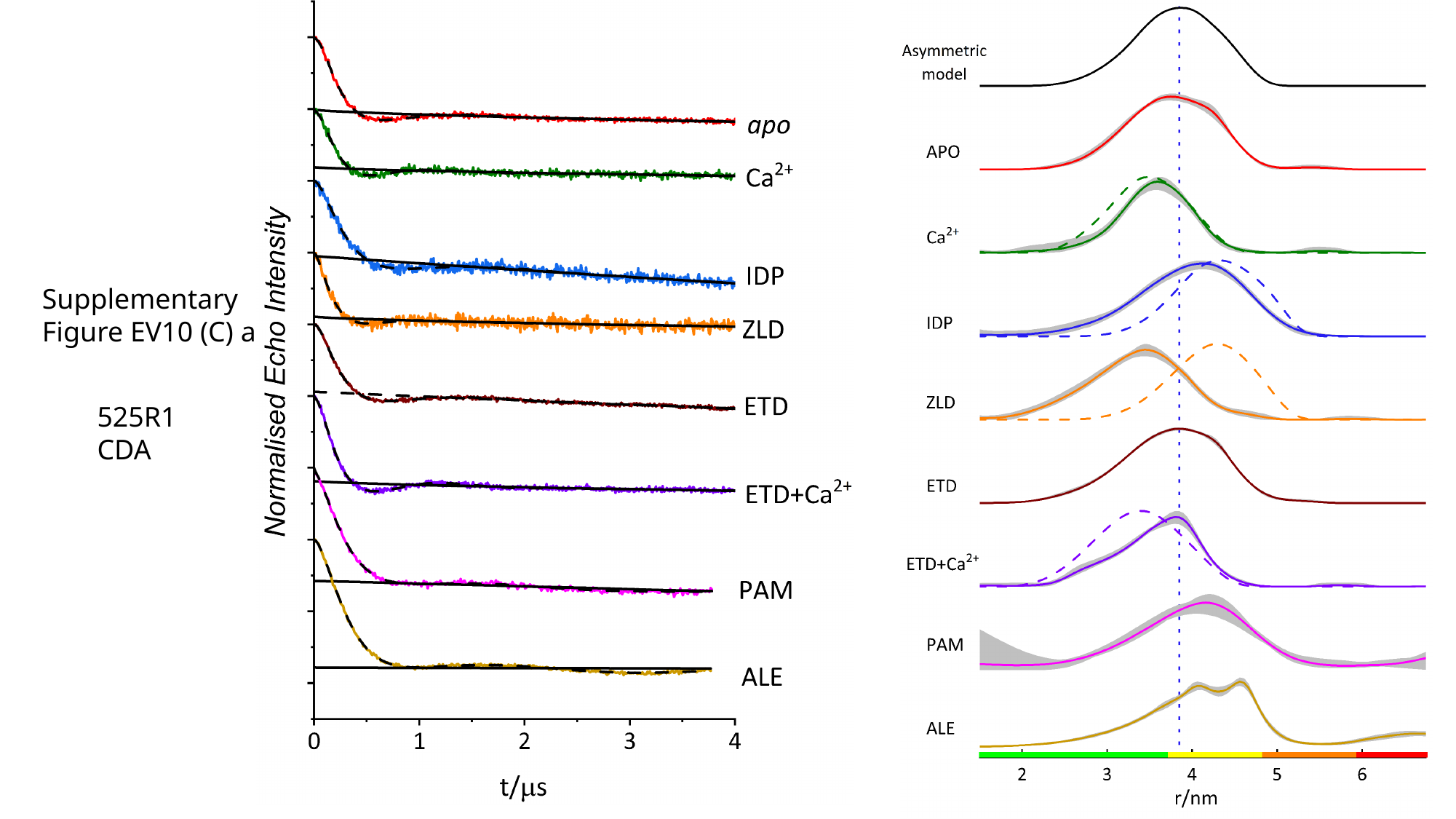

Supplementary
Figure EV10 (C) and (D)
525R1
CDA

### Slide 10
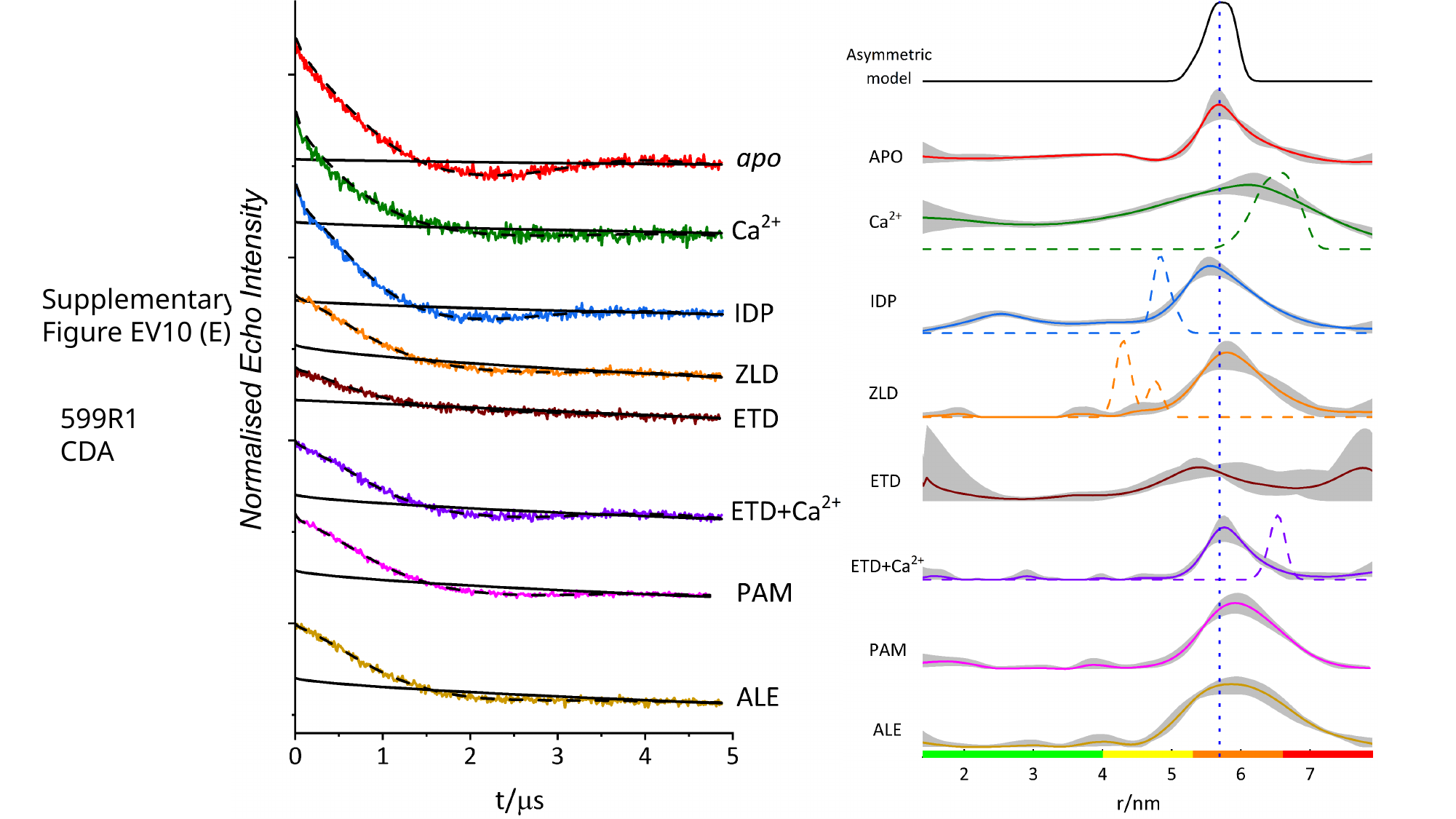

Supplementary
Figure EV10 (E) and (F)
599R1
CDA

### Slide 11
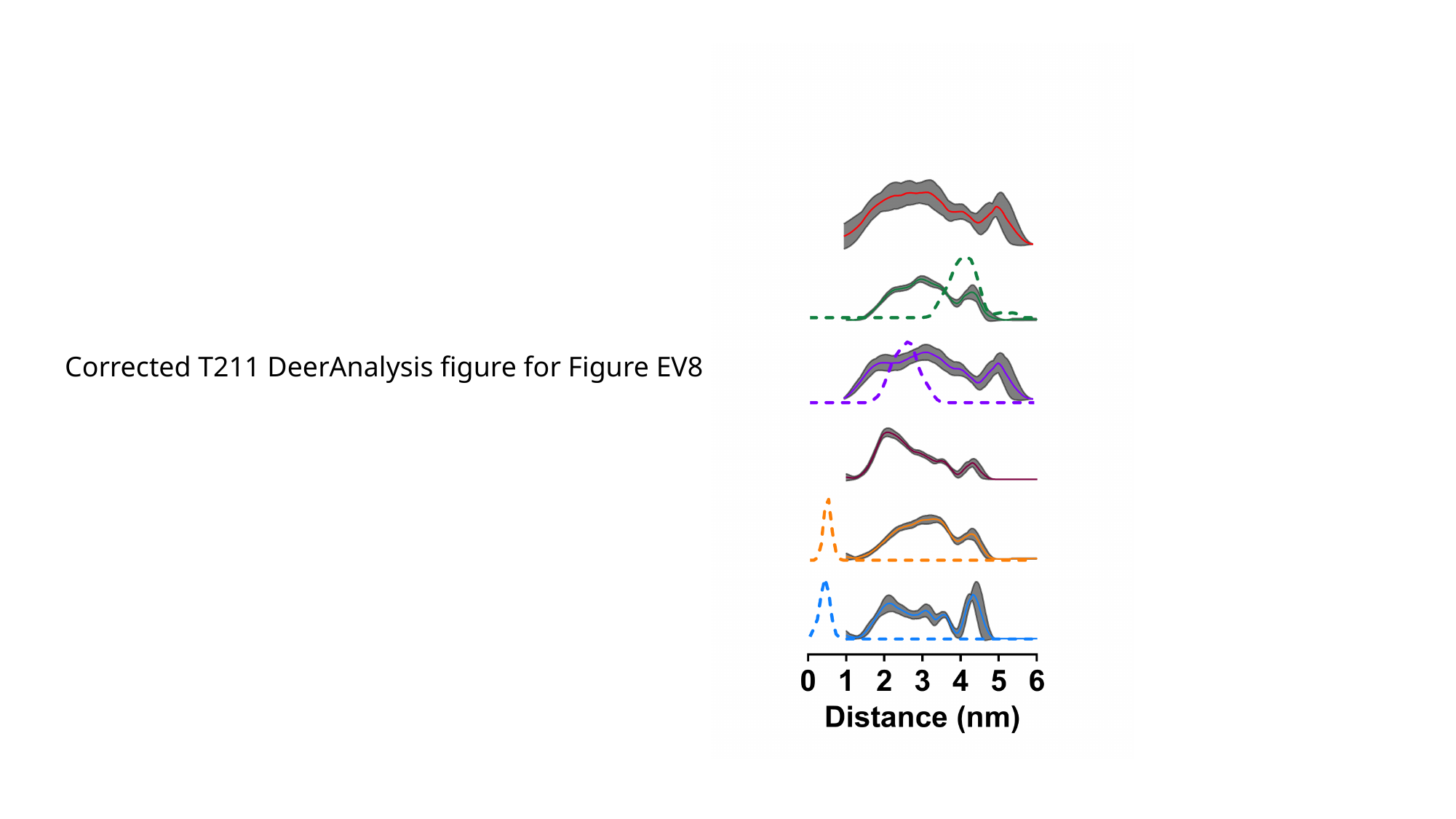

Corrected T211 DeerAnalysis figure for Figure EV8

### Slide 12
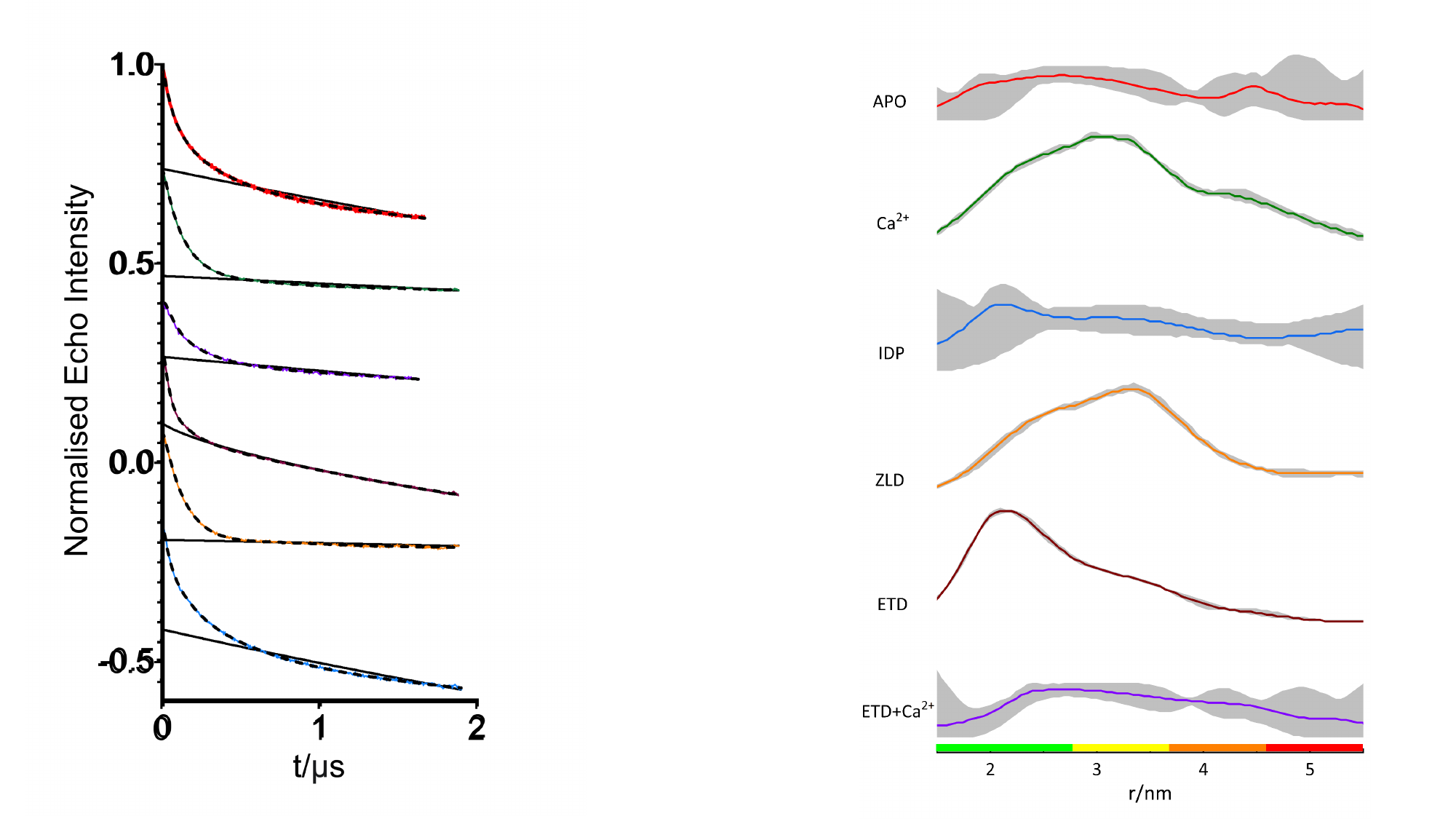

#
